# Xylose Metabolism and the Effect of Oxidative Stress on Lipid and Carotenoid Production in *Rhodotorula toruloides*: Insights for Future Biorefinery

**DOI:** 10.1101/2020.05.28.121012

**Authors:** Marina Julio Pinheiro, Nemailla Bonturi, Isma Belouah, Everson Alves Miranda, Petri-Jaan Lahtvee

## Abstract

The use of cell factories to convert sugars from lignocellulosic biomass into chemicals in which oleochemicals and food additives, such as carotenoids, is essential for the shift towards sustainable processes. *Rhodotorula toruloides* is a yeast that naturally metabolises a wide range of substrates, including lignocellulosic hydrolysates, and converts them into lipids and carotenoids. In this study, xylose, the main component of hemicellulose, was used as the sole substrate for *R. toruloides*, and a detailed physiology characterisation combined with absolute proteomics and genome-scale metabolic models was carried out to understand the regulation of lipid and carotenoid production. To improve these productions, oxidative stress was induced by hydrogen peroxide and light irradiation and further enhanced by adaptive laboratory evolution. Based on the online measurements of growth and CO_2_ excretion, three distinct growth phases were identified during batch cultivations. The intracellular flux estimations correlated well with the measured protein levels and demonstrated improved NADPH regeneration, phosphoketolase activity and reduced beta-oxidation, correlating with increasing lipid yields. Light irradiation resulted in 70% higher carotenoid and 40% higher lipid yields. The presence of hydrogen peroxide did not affect the carotenoid yield but culminated in the highest lipid yield of 0.65 g/g_DCW_. The adapted strain showed improved fitness and 2.3-fold higher carotenoid yield than the parental strain. This work presented a holistic view of xylose conversion into microbial oil and carotenoids by *R. toruloides* for further cost-effective and renewable production of these molecules.

## 1 Introduction

*Rhodotorula toruloides* is considered one of the most promising oleaginous yeasts for industrial applications. This microorganism is a natural producer of lipids (microbial oil) and high-value compounds, such as carotenoids and enzymes for pharma and chemical industries (L-phenylalanine ammonia-lyase and D-amino acid oxidase) (Park et al., 2017). The microbial oil, which is primarily composed of triacylglycerides (TAGs), is a potential raw material for oleochemicals that can be used as biodiesel, cosmetics, and coatings as well as a replacement of vegetable oil in fish feed (Unrean et al., 2017; Blomqvist et al., 2018; Yang et al., 2018). Carotenoids are important molecules for different industries, such as the food, chemical, pharmaceutical and cosmetics industries. In addition to its colorants properties, carotenoids can be metabolised into vitamin A and have antioxidant activity that has been explored, for example, in the prevention of cancer, immune diseases and as skin protection against radiation (Stahl and Sies, 2007; Du et al., 2016; Kot et al., 2019). The global market for carotenoids should reach US$2.0 billion by 2022 (BBC Research, 2018), while the global demand for fatty acids (FAs) and alcohols is expected to reach over 10 Mt in 2020 (Adrio, 2017).

In addition to the ability to produce a variety of relevant compounds, *R. toruloides* can consume a range of carbon and nitrogen sources (Park et al., 2017; Lopes et al., 2020a), including lignocellulosic hydrolysates (Bonturi et al., 2017; Lopes et al., 2020b). Following cellulose, hemicellulose is the second most abundant fraction of lignocellulose, and this fraction consists of polymerised five-carbon sugars, mainly xylose. The efficient utilisation of xylose by a microorganism is essential to improve the conversion of lignocellulosic materials into target compounds, thus increasing the economic viability of the biotechnological processes in biorefineries. Therefore, understanding of the metabolic mechanisms involved in the production of lipids and carotenoids from xylose by *R. toruloides* is crucial to further improve the titres, yields and rates of this bioprocess.

The cellular content of lipids and carotenoids is affected by several factors, including medium composition and cultivation conditions (Mata-Gómez et al., 2014). Previous studies have described the increase in carotenoid production in the presence of oxidative stress, such as hydrogen peroxide (H_2_O_2_) and light irradiation. In the presence of H_2_O_2_ (5 mmol/L), a 5-fold increase in carotenoid production by *Rhodotorula mucilaginosa* was observed (Irazusta et al., 2013), illustrating how optimisation of cultivation conditions can improve production yields of the desired metabolites. Under light irradiation (4,000 lux), the production of carotenoids and lipids by *Rhodotorula glutinis* increased 60% (Gong et al., 2020). The cellular response mechanism against oxidative stress is not clear for yeast, including *R. toruloides.* This condition is associated with the presence of reactive oxygen species (ROS), including H_2_O_2_, superoxide (O_2_^-^) and hydroxyl radicals (HO•). ROS are potent oxidants that can damage all cellular components, including DNA, lipids and proteins. Microbial cells possess two defensive systems against oxidative damage: enzymatic and non-enzymatic. The former is mainly constituted by enzymes superoxide dismutase and catalase, and the latter is involved in direct scavenging of ROS or recycling of oxidised compounds, such as ascorbate, glutathione, alpha-tocopherol and carotenoids (Irazusta et al., 2013).

Integration of large-scale data sets is crucial for a better understanding of the genomic organisation and metabolic pathways in living cells. Complete genome sequences are available for several *R. toruloides* strains (Kumar et al., 2012; Zhu et al., 2012; Morin et al., 2014; Hu and Ji, 2016; Sambles et al., 2017; Coradetti et al., 2018). The lipid formation process during different growth phases of cultivation on glucose has been investigated through proteomic analysis (Liu et al., 2009) and compared with cells grown on xylose (Tiukova et al., 2019b). Multi-omics analyses have identified metabolism modification under nitrogen (Zhu et al., 2012) and phosphate limitation (Wang et al., 2018b). The latter studies have identified higher lipid accumulation under nitrogen or phosphate limitation, which has been correlated to higher activation in nitrogen recycling but also lipid degradation and autophagy. Carotenoid production from glycerol was investigated using global metabolomics, revealing reduced abundance of metabolites involved in TCA and amino acid biosynthesis (Lee et al., 2014).

Genome-scale metabolic models (GEMs) are another powerful tool to understand and provide a holistic view of metabolic fluxes, energy and redox metabolism or even suggest targets for metabolic engineering. GEMs are constructed based on the available genome sequence of a specific organism, thus providing a summary of the metabolic network (Kerkhoven et al., 2015). Regarding *R. toruloides*, a number of metabolic models are available to assess lipid production (Bommareddy et al., 2015; Castañeda et al., 2018), and two reports of GEMs are available (Dinh et al., 2019; Tiukova et al., 2019a). Lopes et al. (2020a) reported the first study that combined data from cultivations of *R. toruloides* under different carbon sources with the GEM. The approach proved to be useful for understanding metabolic fluxes and identifying targets to improve lipid production either by metabolic engineering or process optimisation.

Therefore, the current study aimed at providing a holistic view of lipid and carotenoid production by *R. toruloides* using xylose as a sole carbon source by combining detailed physiological characterisation with the quantitative proteomics and GEM analysis. Oxidative stress (H_2_O_2_ and light irradiation) and adaptive laboratory evolution were employed to improve lipid and carotenoids production. To our knowledge, this is the first work combining such approaches for this strain, and the data obtained here can be used to establish future bioprocesses in biorefineries.

## 2 Material and methods

### 2.1 Strain and inoculum

*Rhodotorula toruloides* CCT 7815 was obtained from “Coleção de Culturas Tropicais” (Fundação André Tosello, Campinas, Brazil) and stored at −80°C in 10% (v/v) glycerol. This strain was derived from *R. toruloides* CCT 0783 (synonym IFO10076) after short-term adaptation in sugarcane bagasse hemicellulosic hydrolysate (Bonturi et al., 2017). The inoculum was prepared in YPD medium at 200 rpm and 30°C for 24 h. The cells were washed twice with 0.9% (v/v) NaCl before inoculation.

### 2.2 Adaptive laboratory evolution

Adaptive laboratory evolution (ALE) was performed by successive shake flask cultivations of *R. toruloides* in the presence of H_2_O_2_ in rich medium (30.0 g xylose, 2.5 g glucose, 0.9 g yeast extract, 0.2 g (NH_4_)2SO_4_, 1.5 g KH_2_PO_4_ and 0.9 g MgSO_4_·7 H_2_O). ALE was performed in two cycles with the aim of improving the performance of this yeast under this oxidative environment. The initial H_2_O_2_ concentration was 10 mmol/L (the highest concentration in which the cells grew in successive cultivations) in the first cycle and 20 mmol/L in the second cycle (the highest concentration that the parental strain tolerated). In each cycle, the cells were harvested at the exponential growth phase and transferred to a fresh medium with the same H_2_O_2_ concentration. The initial OD (at 600 nm) of every passage was 0.5. The end of the cycle was determined by the stabilisation of the maximum specific growth rate and the length of lag phase.

### 2.3 Yeast cultivation

Batch cultivations were performed in 1-L bioreactors (Applikon Biotechnology, Delft, The Netherlands) with a working volume of 600 mL at pH 6.0 and controlled with the addition of 2 mol/L KOH. Dissolved oxygen was maintained at greater than 25% using 1-vvm airflow and stirring speeds between 400 and 600 rpm. The cultivation started with 1% (v/v) of overnight culture inoculum. Oxidative stress was induced by the addition of H_2_O_2_ (20 mmol/L) at the start of the cultivation or by light irradiation (white LED light, 40,000 lux) throughout the experiment. The composition of CO_2_ and O_2_ in the gas outflow was measured using an online gas analyser (BlueSens GmbH, Herten, Germany), and biomass was monitored online using a Bug Lab BE3000 Biomass Monitor (Bug Lab, Concord, CA, USA) at 1,300 nm. Data were collected and processed with BioXpert V2 software v. 2.95 (Applikon Biotechnology, Delft, the Netherlands). All experiments were performed in triplicate.

The medium composition used in the bioreactor experiments was, per litre, 70.0 g xylose, 1.95 g (NH_4_)2SO_4_, 3.0 g KH_2_PO_4_, 0.5 g MgSO_4_·7 H_2_O, 1.0 mL vitamins solution and 1.0 mL trace metal solution (Lahtvee et al., 2017), supplemented with 100 μl antifoam 204 (Sigma-Aldrich, MO, USA). Samples for biomass, carotenoid and extracellular metabolite analyses were collected every 24 h. Samples were withdrawn from bioreactors, transferred into precooled 2-mL Eppendorf tubes and centrifuged for 20 s at 4°C and 18,000 g. The supernatant was collected and stored at −20°C for extracellular metabolite quantification. The biomass was snap-frozen in liquid nitrogen, stored at −80°C, and further used for proteomics analysis.

### 2.4 Quantification of biomass, extracellular metabolites, carotenoids, total lipids and proteins

Online turbidity measurements were calibrated by gravimetrically measuring the dry cellular weight (DCW) every 24 h. Extracellular metabolites in the broth were measured using HPLC (LC-2030C Plus, Shimadzu, Kyoto, Japan) equipped with a refractive index detector (RID-20A, Shimadzu, Kyoto, Japan). Concentrations of xylose, organic acids and glycerol were measured using an HPX-87H column (Bio-Rad, CA, USA) at 45°C, and 5 mmol/L H_2_SO_4_ served as the mobile phase with isocratic elution at 0.6 mL/min. Xylitol and arabitol were quantified using Rezex RPM-Monosaccharide column (Phenomenex, CA, USA) at 85°C, and purified water (MilliQ Ultrapore Water System, Merck, Darmstadt, Germany) used as the mobile phase with isocratic elution at 0.6 mL/min.

For quantification of carotenoids (modified from Lee et al., 2014), 2 mL of cells were harvested by centrifugation, washed twice in distilled water and resuspended in 1.0 mL of acetone. The cells were lysed with acid-washed glass beads (400 – 650 μm) using the FastPrep homogeniser for three cycles (4 m/s for 20 s) (MP Biomedicals, CA, USA). After centrifugation at 15,000 g for 5 min, the acetone solution containing carotenoids was collected and stored at 4°C. These steps were repeated until the cell debris was colourless. Then, the solvent was evaporated in Concentrator Plus (Eppendorf, Hamburg, Germany), and the remaining extracts were resuspended in a known volume of acetone. Carotenoids were measured using Acquity UPLC (Waters, MA, USA) equipped with a TUV detector (Waters, MA, USA) and C18 column (BEH130, 1.7 μm, 2.1 x 100 mm, Waters, MA, USA). The mobile phase was a gradient from 80 to 100% of acetone in purified water at a flow rate of 0.2 mL/min. Detection was performed at 450 nm (modified from Weber et al., 2007). All identified peaks were quantified using the β-carotene standard (Alfa Aesar, MA, USA). Detected peaks were identified according to the known carotenoid retention time profile (Weber et al., 2007; Lee et al., 2014).

Lipids were extracted according to an adaptation of the Folch method (Folch et al., 1957) described by Bonturi et al. (2015). Briefly, a mixture of chloroform and methanol (2:1 v/v) was added to dried biomass. After 24 h, the solvent was evaporated in a rotary evaporator (Buchi, Switzerland), and the total lipid content was determined gravimetrically.

Total proteins were extracted from 600 μg of biomass resuspended in Y-PER solution (Thermo Fisher, MA, USA) in a 2-mL Eppendorf tube. This suspension was incubated at 30°C for 45 min. Then, glass beads were added in the tube, and cell lysis was performed in a FastPrep homogeniser during ten cycles (4 m/s for 20 s). Between the cycles, the tubes were placed on ice for 3 min. After centrifugation at 18,000 g and 4°C for 10 min, the supernatant was carefully removed and stored at 4°C for further protein quantification. Y-PER reagent was added to the remaining biomass in the tube, and the cell disruption process was repeated. This step was performed until no protein was detected. Total protein was quantified using a commercially available assay (Micro BCA™ Protein Assay Kit, Thermo Fisher), and samples were diluted in the linear range of BSA protein standard (0.5 to 20 μg/ml).

### 2.5 Proteome analysis

Fully labelled biomass to be used as the internal standard in absolute proteome analysis was produced by cultivating *R. toruloides* in minimal mineral medium containing labelled heavy 15N, 13C-lysine (Silantes, Munich, Germany). Heavy labelling of the proteinogenic lysine was measured at 96.6% (data not shown). Absolute proteomics and internal heavy-labelled standard preparation and analyses were performed similarly as described in Lahtvee et al. (2017) and Kumar and Lahtvee (2020). Briefly, cells were resuspended in the lysis buffer (6 mol/L guanidine HCl, 100 mmol/L Tris-HCl pH 8.5 and 100 mmol/L dithiothreitol) and homogenised with glass beads using the FastPrep24 device. The supernatant was removed by centrifugation (17,000 g for 10 min at 4°C) and precipitated overnight with 10% trichloroacetic acid (TCA) at 4°C. Pellets from the previous precipitation step were spiked in 1:1 ratio with the heavy-labelled standard. This mixture was further precipitated with 10% TCA. The pellet was resuspended in a buffer containing 7 mol/L urea and 2 mol/L thiourea in 100 mmol/L ammonium bicarbonate (ABC) followed by reduction using 5 mmol/L DTT and alkylation with 10 mmol/L chloroacetamide. Peptides were digested at room temperature with *Achromobacter lyticus* Lys-C (Wako Pure Chemical Industries, Osaka, Japan) for 4 h at the ratio of 1:50 (enzyme:protein) followed by overnight digestion of the previous solution diluted 5 times in 100 mmol/L ABC buffer. Peptides were desalted using in-house prepared C18 (3M Empore, Maplewood, MO, USA) tips and were reconstituted in 0.5% trifluoroacetic acid (TFA). For separation, 2 μg of peptides was injected on an Ultimate 3000 RSLCnano system (Dionex, Sunnyvale, CA, USA) coupled to a C18 cartridge trap-column in a backflush configuration and an analytical 50 cm x 75 μm emitter-column (New Objective, MA, USA) in-house packed with 3 μm C18 particles (Dr. Maisch, Ammerbuch, Germany). Eluted peptides were sprayed to a quadrupole-orbitrap Q Exactive Plus (Thermo Fisher Scientific, MA, USA) tandem mass spectrometer. MaxQuant 1.4.0.8 software package was used for raw data identification and identification (Cox and Mann, 2008). *R. toruloides* NP11 served as a reference proteome database in UniProt (www.uniprot.org). Protein quantification was performed following the total protein approach described in (Sánchez et al., 2020) and assuming 90% coverage from the total protein abundance.

LC-MS/MS proteomics data were deposited in the ProteomeXchange Consortium (http://proteomecentral.proteomexchange.org) via the PRIDE partner repository (Vizcaı’no et al., 2013) and can be retrieved using the dataset identifier PRIDE: PXD019305. Processed quantitative data are presented in Supplementary Table S7. Triplicated quantitative proteomics data were used for differential expression analysis. P-values were adjusted for multiple testing using the Benjamini-Hochberg procedure (Benjamini and Hockberg, 1995). Additional data analysis included gene set enrichment analysis (carried out using PIANO platform; Väremo et al., 2013) and gene enrichment analysis (g:Profiler; Raudvere et al., 2019).

### 2.6 Genome-scale modelling

The intracellular flux patterns were predicted using version 1.2.1. of the *R. toruloides* metabolic network at the genome scale (Tiukova et al., 2019a; Lopes et al., 2020a) and rhto-GEM (https://github.com/SysBioChalmers/rhto-GEM/releases/). The model was improved by adding carotenoids (β-carotene, γ-carotene, torulene and torularhodin) into the biomass reaction. Flux balance analysis was performed to calculate internal fluxes using RAVEN Toolbox (Wang et al., 2018a) on MATLAB (The MathWorks Inc., MA, USA), Gurobi solver (Gurobi Optimization Inc., TX, USA) and by optimising for non-growth related ATP maintenance (NGAM). The latter was followed by flux variability analysis (random sampling at n = 5000) at 95% from the maximal NGAM value. Experimental data obtained from the yeast cultivations were used to constrain the model if not stated otherwise. Biomass composition was adjusted to the measured total protein, lipid and carotenoid content.

## 3 Results

### 3.1 Three distinct phases of *R. toruloides* growth on xylose

*R. toruloides* growth was characterised with xylose as a sole carbon source under aerobic batch conditions on a mineral medium. The initial xylose concentration of 70 g/L was chosen, and the amount of nitrogen was adjusted to result in a C/N ratio of 80 mol/mol. On-line monitoring of culture turbidity, CO_2_ production and O_2_ consumption was complemented by off-line analysis of sugars, alcohols and biomass composition (total lipids, proteins and carotenoids; Figure 1). During the batch cultivation of *R. toruloides*, three distinct growth phases were observed based on growth dynamics and substrate consumption patterns. In the first growth phase (P1), cells were growing exponentially without any observable limitation, and xylose used as the sole carbon source. Arabitol, xylitol and CO_2_ were the main fermentation by-products detected. Phase two (P2) started with a sudden decrease in the specific growth rate due to nitrogen limitation. At that point, approximately 23 g/L of xylose was consumed, indicating a critical C/N ratio of 26 for *R. toruloides* to reach nitrogen limitation. P2 lasted until the depletion of the primary carbon source, namely, xylose. Consumption of arabitol and xylitol under nitrogen limitation defined the third growth phase (P3). To our knowledge, no previous study has provided a characterisation of *R. toruloides* physiology on xylose in such detail. These phases were further analysed in this work, aiming to identify the cellular metabolic behaviour in response to the environment changes during the batch growth.

**Figure 1.**
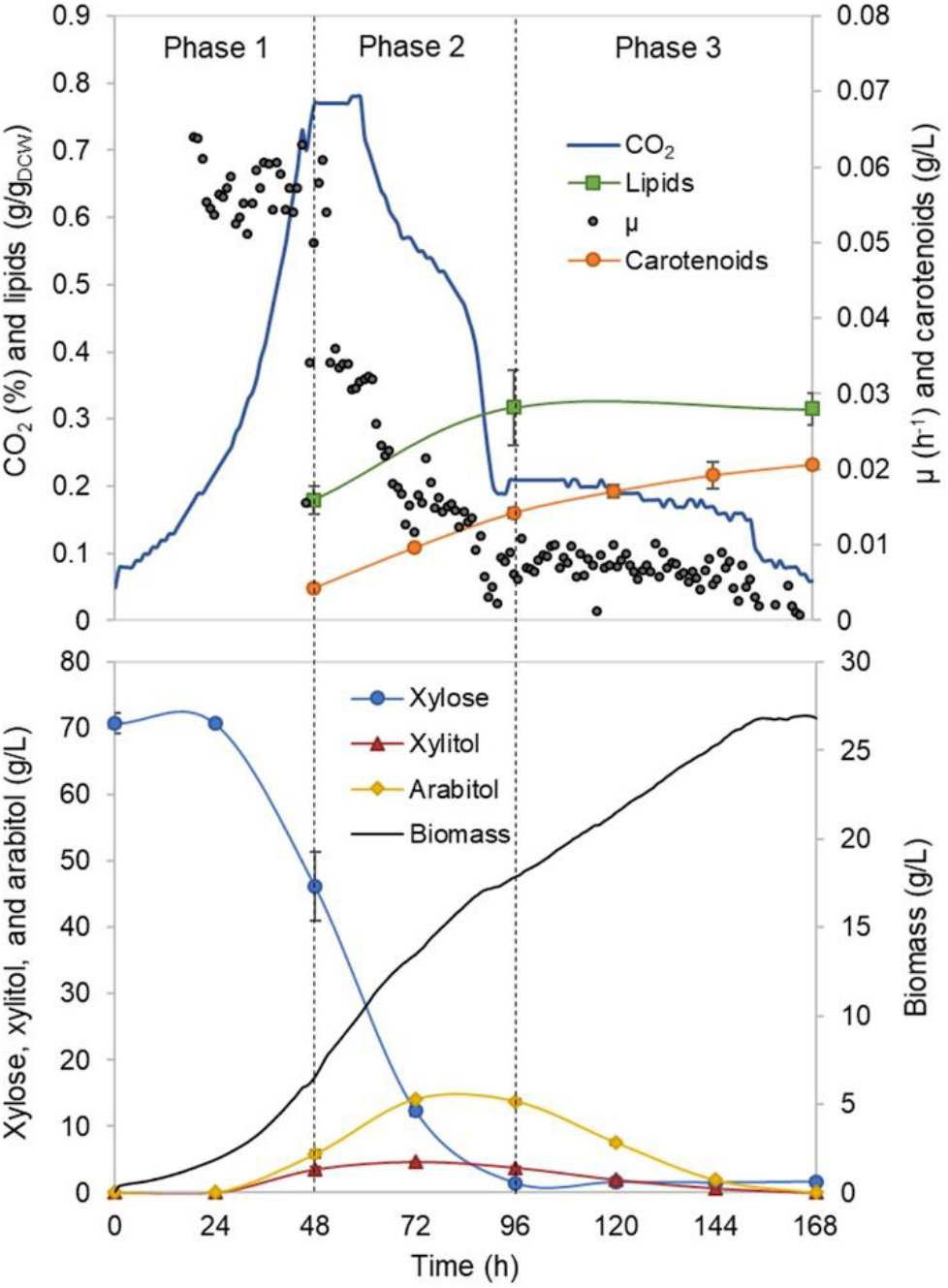
*R. toruloides* batch cultivation on xylose under the reference (optimal) environmental conditions for the parental strain. Dashed vertical lines define three observed growth phases. The specific growth rate (μ), biomass concentration, intracellularly accumulated lipid and carotenoid concentrations, extracellular metabolite profiles and CO_2_ production profile in the outflow gas are presented. The values represent an average of three independent cultivation experiments; error bars represent standard deviation.

P1 comprised the highest specific growth rate (μ, 0.060 ± 0.001 h^−1^) and the specific xylose uptake rate (r_XYL_, 1.74 ± 0.12 mmol/g_DCW_.h) (Supplementary Table S1), while no nutrient-level limitations was detected. During the exponential growth, approximately one-third of the consumed carbon was secreted as arabitol and xylitol. An additional one-third of the consumed carbon was secreted as CO_2_. Although the carotenoid yield in P1 was low (0.66 ± 0.06 mg/g_DCW_), the specific production rate (r_CAR_) was the highest (0.042 ± 0.003 mg/g_DCW_.h) due to the higher μ (Figure 2A, Supplementary Table S1). The biomass and lipid yields were 0.25 ± 0.02 g_DCW_/g and 0.18 ± 0.03 g/g_DCW_, respectively. Carbon balance in this phase was estimated at 93%, indicating a small amount of undetected by-products.

**Figure 2.**
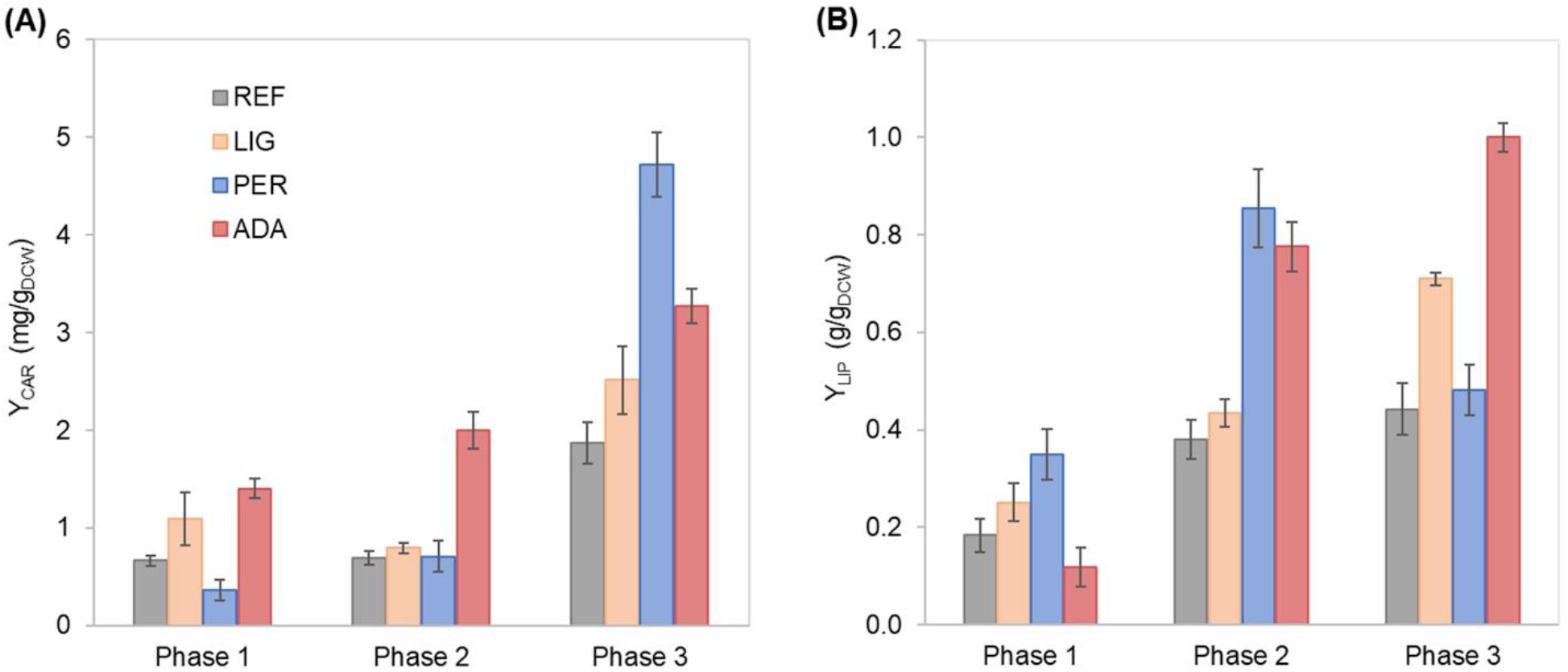
Carotenoid **(A)** and lipid **(B)** yields on biomass in each growth phase of *R. toruloides* parental strain under the reference condition (REF), light irradiation (LIG) and in the presence of hydrogen peroxide (PER), and the adapted strain under oxidative stress (ADA). Yields were calculated considering the production in each growth phase.

Nitrogen limitation under the xylose-excess conditions marked the start of P2. The limitation led to a decrease in μ (average value 0.020 ± 0.002 h^−1^) and r_XYL_ (average value 0.41 ± 0.06 mmol/g_DCW_.h). Compared to P1, the carotenoid yield did not show a significant difference; however, r_CAR_ decreased 4-fold compared with P1 (Supplementary Table S1). As expected, lipid accumulation doubled compared to P1 due to the positive influence of nitrogen limitation, reaching 0.38 ± 0.05 g/g_DCW_ (Figure 2B). P3 started after the depletion of xylose. Here, xylitol and arabitol were simultaneously consumed by the cells. The average specific growth rate was the lowest of the three growth phases (0.005 ± 0.0003 h^−1^). The highest accumulation of carotenoids per biomass was detected in this phase, increasing substantially to 1.87 ± 0.21 mg/g_DCW_; however, the r_CAR_ was the lowest of all phases (Figure 2 and Supplementary Table S1). The lipid yield remained at the same level as that noted in P2. In addition, 50% carbon loss (undetected carbon) was observed in P3, which can be partially explained by the technical uncertainty of measurements for off-gas at the very low growth rate conditions in the P3. Total carotenoid and lipid yields for the whole batch cultivation were 0.85 ± 0.01 mg/g_DCW_ and 0.33 ± 0.07 g/g_DCW_, respectively (Table 1, column REF).

### 3.2 Understanding intracellular flux patterns among three growth phases

To understand the changes in intracellular flux patterns between the observed growth phases, simulations using *R. toruloides* GEM were performed (Supplementary Tables S2; Tiukova et al., 2019a; Lopes et al., 2020a). GEM is a mathematical reconstruction of the metabolic network based on genome annotations and information, such as gene-protein-reaction relationships. GEM in combination with flux balance analysis (FBA) allows simulation of intracellular flux patterns. In addition to intracellular rates, FBA can be used to predict potential by-products, which were not identified experimentally. Given that 7 and 5% of carbon was missing in P1 and P2, respectively, the model predicted that glyoxylate and acetaldehyde as the most likely undetected products secreted in P1 and P2, respectively. For the flux distribution comparison between the observed growth phases, fluxes were normalised by the total carbon uptake rate (Supplementary Tables S3).

NADPH supply is crucial for lipid production given that every elongation step of fatty acid (FA) synthesis requires the oxidation of two NADPH (Wasylenko et al., 2015). In addition to FA synthesis, xylose and L-arabitol utilization also require NADPH. For all three growth phases, the highest NADPH usage was observed for substrate uptake: xylose reductase (r_1093) in P1 and P2 and L-xylulose reductase (t_0882) in P3 (Supplementary Figure S1, Table S5). During xylose metabolism (P1 and P2), arabitol production via L-xylulose reductase partially regenerated the oxidised NADPH. Once xylose was exhausted (P3), NADPH was required for arabitol catabolism (Supplementary Figure S1). Our simulations noted that the oxidative branch of the pentose phosphate pathway (PPP), namely, glucose 6-phosphate dehydrogenase (r_0466) and phosphogluconate dehydrogenase (r_0889), was responsible for 83, 87 and 96% of NADPH regeneration in P1, P2 and P3, respectively. Although NADPH demand in the substrate consumption and amino acid biosynthesis pathways decreased in P2 and P3 compared to P1, fluxes in lipid biosynthesis increased 1.4-fold.

At the xylulose-5P branch point, 91% of carbon entered into the central carbon metabolism via traketolase (r_1049, r_1050) in P1 (Supplementary Figure S1). The remaining 9% was converted into glyceraldehyde-3-phosphate and acetyl-phosphate by the phosphoketolase reaction (t_0081). Under nitrogen limitation, the activity of phosphoketolase was approximately tripled compared to P1. The phosphoketolase pathway further generates acetyl-CoA, a precursor of FA synthesis, by phosphate transacetylase (t_0082) without losing a carbon compared to the pathway originating from pyruvate.

At the pyruvate branch point, on average, ca. 70% of the pyruvate produced in the cytosol was transported to mitochondria (r_1138 and r_2034) to be converted by pyruvate dehydrogenase (r_0961) into acetyl-CoA, which is used in the tricarboxylic acid (TCA) cycle by citrate synthase (r_0300). The remaining cytosolic pyruvate was either converted to cytosolic oxaloacetate (pyruvate carboxylase, r_0958) or into acetyl-CoA by three enzymatic steps (pyruvate decarboxylase, r_0959; acetaldehyde dehydrogenase, r_2116; acetyl-CoA synthase, r_0112). Acetyl-CoA can also be produced from citrate by ATP citrate lyase (y200003) or from acetyl-P by phosphate transacetylase. Under excess nitrogen conditions (P1), acetyl-CoA synthase was responsible for 60% of the flux. However, under nitrogen limitation (P2 and P3), ca. 71% of acetyl-CoA originated via phosphate transacetylase (Supplementary Figure S1).

Our physiological data showed that the lipid yield was higher under nitrogen-limitation phases (P2 and P3), and the carotenoids yield was higher in P3. These results can be explained by the higher predicted fluxes through reactions involving phosphoketolase, FA and carotenoids synthesis, and NADPH regeneration.

### 3.3 Impact of oxidative stress via light irradiation or the presence of H_2_O_2_

Further, we were interested in how oxidative stress created by either 20 mmol/L H_2_O_2_ or light irradiation (40,000 lux white light) affects cellular growth and lipid and carotenoid accumulation. Cultivation of *R. toruloides* under these oxidative stress conditions presented the same three growth phases described for the reference condition (REF) and a similar growth and substrate consumption profile (Supplementary Figure S2 A, B). The specific growth rate differences compared to the reference condition were insignificant under the light irradiation (LIG) condition, but a significant 50% decrease was observed under the H_2_O_2_ stress (PER) in P1 (Supplementary Table S1). The most significant difference was detected in the longer lag phase (approximately 90 h) shown in the PER condition (Supplementary Figure S2B). Moreover, the accumulation profiles of carotenoids and lipids showed altered behaviour compared to the reference (Figure 2). The stress caused by H_2_O_2_ negatively affected carotenoid production in P1. However, in P3, the total carotenoid yield was the highest of all conditions (4.72 ± 0.47 mg/g_DCW_); thus, the highest r_CAR_ (0.026 ± 0.012 mg/g_DCW_.h) was reached in the third phase (Figure 2A, Supplementary Table S1). Lipid production exhibited a different behaviour, presenting a higher yield in P1 and P2 compared to the reference condition. The specific lipid production rate (rLIP) in P2 was the highest of all conditions in all phases (0.020 ± 0.004 g/g_DCW_.h), albeit the specific growth rate was not amongst the highest obtained in this study.

Regarding the overall results, for the entire cultivation under light irradiation, the cells presented 70% increased carotenoid yield (1.45 ± 0.14 mg/g_DCW_) and 40% increased lipid yield (0.46 ± 0.12 g/g_DCW_) compared to the reference condition. H_2_O_2_ stress did not affect carotenoid production by the parental strain, and the production yield was maintained at 0.89 ± 0.04 mg/g_DCW_. Surprisingly, this condition showed the highest lipid yield (0.65 ± 0.06 g/g_DCW_), which was increased by two-fold compared with the reference condition (Table 1). The achieved lipid yield was only slightly lower than the highest lipid yield reported in the literature for *R. toruloides;* specifically, 0.68 g/g_DCW_ has been reported in fed-batch cultivation on a rich glucose-based medium (Li et al., 2007).

**Table 1.** Global titres, yields on biomass (Y), and volumetric production rate (q) of carotenoids and lipid production by *R. toruloides* parental strain under the reference condition (REF), light irradiation (LIG), hydrogen peroxide stress (PER), and the adapted strain under oxidative stress (ADA). These values were based on the final titres and the total fermentation time, representing the mean and standard deviation of triplicate experiments.

|  |  | REF | LIG | PER | ADA |
| --- | --- | --- | --- | --- | --- |
| Carotenoids | Titre (mg/L) | 20.7 ± 0.5 | 36.2 ± 2.6 | 19.3 ± 0.2 | 44.0 ± 2.4 |
| | $Y_{CAR}$ (mg/DCW) | 0.85 ± 0.01 | 1.45 ± 0.14 | 0.89 ± 0.04 | 1.90 ± 0.13 |
| | $q_{CAR}$ (mg/L.h) | 0.12 ± 0.003 | 0.22 ± 0.02 | 0.06 ± 0.02 | 0.23 ± 0.01 |
| Lipids | Titre (mg/L) | 8.1 ± 0.3 | 11.1 ± 1.7 | 13.8 ± 0.9 | 13.3 ± 0.5 |
| | $Y_{LIP}$ (g/g<sub>DCW</sub>) | 0.33 ± 0.07 | 0.46 ± 0.12 | 0.65 ± 0.06 | 0.58 ± 0.07 |
| | $q_{LIP}$ (g/L.h) | 0.05 ± 0.003 | 0.07 ± 0.005 | 0.05 ± 0.002 | 0.07 ± 0.002 |

### 3.4 Increasing carotenoid accumulation via adaptive laboratory evolution

Adaptive laboratory evolution (ALE) is a strategy to improve the fitness of the microorganism in a challenging environment. H_2_O_2_ stress improved lipid accumulation but increased the lag phase and lowered the growth rate. Therefore, ALE was performed to improve those parameters. The first cycle of ALE started with 10 mmol/L of H_2_O_2_. After 16 passages (ca. 30 generations), the lag phase decreased 11-fold (from 46 h to 4 h), and μmax stabilised at 0.045 ± 0.003 h^−1^, resulting in a 66% increase compared to the parental strain under the same conditions (Supplementary Figure S3A). Once the μmax plateaued, the second cycle of ALE was started by increasing the selective pressure to 20 mmol/L of H_2_O_2_ in the medium (Supplementary Figure S3B). As a result, after 15 passages (ca. 20 generations), the lag phase of yeast growth decreased from 30 to 5 h, and the μmax improved to 0.055 ± 0.001 h^−1^, representing a 22% increase compared to the first cycle. Although μmax did not improve remarkably during the second cycle of ALE, the cells presented stronger pink colouration compared to the parental strain, indicating increased carotenoid accumulation. Therefore, the ALE experiment was halted, and the strains were characterised.

The adapted strain in presence of H_2_O_2_ (20 mmol/L) showed a remarkable ca. 3-fold increase in carotenoid yield and titre, respectively, compared to the parental strain under the same condition during the initial screening experiments (Supplementary Table S6). Therefore, the adapted strain was further characterised under controlled environmental conditions in bioreactors and studied on the proteomics level.

### 3.5 Adapted strain under H_2_O_2_ stress

Under H_2_O_2_ stress, the adapted strain (ADA) exhibited a 70-hour shorter lag phase compared with the parental strain (PER) (Supplementary Figure S2C). Aeration and agitation in the bioreactor may increase the oxidative stress, which could explain the longer lag phase compared to initial screening experiments in shake flasks (mentioned above).

Some differences were noticed between the ADA and other conditions. Although the μ in P1 did not show a significant difference, the r_CAR_ was increased by 4-fold in ADA compared with PER (Supplementary Table S1). However, in P2, the μ in ADA was 2-fold reduced compared with the other conditions, but r_CAR_ remained 20% increased. The lower r_XYL_ in P2 could be related to the lower production of xylitol and arabitol in ADA compared to the other conditions probably due to a softer redox imbalance during xylose catabolism. The ADA showed ca. 2-fold increased yield of carotenoids and lipids under nitrogen-limiting phases (P2 and P3) compared to REF. In P3, the lipid yield was 1.0 ± 0.03 g/g_DCW_, indicating that the gain of cell mass noted during this phase was mainly related to lipid accumulation.

The whole batch growth of ADA exhibited a 2.3-fold increase in carotenoid yield compared to PER; however, the lipid yield did not show a significant difference (Table 1).

### 3.6 Composition of carotenoids in biomass

*R. toruloides* mainly produces four carotenoids: γ-carotene, β-carotene, torulene and torularhodin (Mata-Gómez et al., 2014). The carotenoid profile was very similar in all the studied conditions. The β-carotene fraction decreased over time, whereas the opposite was observed for torulene (Supplementary Figure S3). Torularhodin was the most abundant fraction of carotenoids under all studied conditions. Growth of the parental strain under light irradiation and adapted strain cultivations showed torularhodin fraction higher than 50% during P2, which can be related to a stronger antioxidative property of this carotenoid, attributed to the presence of more double bonds in its chemical structure (Kot et al., 2018).

### 3.7 Proteomic results revealed the highest difference between nitrogen excess and -limiting conditions

Total protein measurements were combined with the absolute proteome analysis for the most relevant conditions. Therefore, samples from all three different growth phases from reference cultivation were analysed together with P2 of light-induced oxidative stress (LIG P2) and P1 and P3 of H_2_O_2_-induced oxidative stress for the adapted strain (ADA P1 and P3, respectively). At the latter three sampling points, the production of carotenoids and lipids was increased compared to the REF. One-third increased total protein content was measured for the nitrogen-excess condition during the P1 of a reference culture. All other conditions showed no significant differences in protein content with an average of 0.14 g/g_DCW_ (Figure 4A). In differential expression analysis, proteome data were normalised to a constant protein mass, representing allocation differences for the individual proteins.

On average, more than 3,000 individual proteins were quantified under every condition studied (Supplementary Table S7). Principal component (PC) analysis clearly identified the biggest differences in the data set, which were determined by the switch into nitrogen limitation as indicated by the clear separation of the samples on the first PC, characterising 39% of the changes (Figure 4B). The second PC separated samples based on the use of the adapted strain under the oxidative stress environment (17% of the difference in the data). Altogether, 1,518 proteins showed significant (adj. p-value < 0.01) allocation changes under at least one of the environmental conditions (Figure 4C). Interestingly, no significant allocation (nor abundance) change was detected between the samples collected from reference or light-induced stress conditions under nitrogen limitation; however, significant differences in biomass composition were detected.

### 3.8 Translation and NADPH metabolism were most affected under nitrogen limitation

To understand the main differences in the dataset, gene set enrichment analysis (GSA) was used to identify classes of proteins that are significantly over-represented among the measured proteins and may have an association with a specific phenotype. A variation of GSA-based analysis was conducted. First, protein-Gene Ontology (GO) group relations were received from UniProt database. Second, protein-subsystem relationships were obtained from rhtoGEM, providing more specific information on various metabolic pathways present in *R. toruloides.* Given that PCA divided samples into four separate quadrants based on nitrogen availability and oxidative stress, we focussed on the comparison of these sample clusters throughout the study. Using UniProt-provided GO groups in GSA, 14 groups exhibited significant over-representation with an adj. p-value < 0.01 under the studied conditions (Supplementary Table S8). Most of these groups were related to protein translation, which were downregulated under nitrogen limitation and correlated with the lower specific growth rates under these conditions. A clear correlation between ribosome abundances and specific growth rate has been demonstrated previously for other organisms (Scott et al., 2014; Metzl-raz et al., 2017). When subsystems from rhtoGEM were considered, significant upregulation was detected among carbon metabolism and its subgroups (glycolysis, gluconeogenesis, TCA cycle, glyoxylate and dicarboxylate metabolism; Supplementary Table S9). Interestingly, only the parental strain (under nitrogen-limiting conditions) showed overexpression in fatty acid degradation pathways and downregulation in amino acid biosynthesis pathways (with the exception of the tryptophan pathway, which was upregulated). Differences in the regulation of fatty acid degradation pathways could be responsible for the significantly increased lipid accumulation under the oxidative stress condition.

Based on the enzyme-metabolite relationships present in the rhtoGEM, reporter metabolites were analysed as the third variation of GSA, illustrating metabolites showing significant alterations among enzymes they interact with (Supplementary Table S10). When samples under nitrogen-limitation were compared to samples cultured under excess nitrogen, the most significant upregulation was detected among proteins in proximity to NAD^+^/NADH (Figure 4D). More than 65% of the proteins associated with NAD^+^/NADH showed increased allocation under nitrogen limitation (REF P2 and P3, LIG P2, ADA P3). In contrast, protein allocation decreased significantly for proteins related to NADP^+^/NADPH metabolism. Downregulation was predominantly noted in amino acid biosynthesis pathways, while NADPH consumption in lipid metabolism and the glutamate production pathway showed upregulation. Upregulation of enzymes in glutamate biosynthesis in response to nitrogen starvation have been demonstrated previously (Zhu et al., 2012; Tiukova et al., 2019b). Additionally, metabolites related to lipid synthesis (CoA, acetyl-CoA, acetate, and pyruvate) showed increased protein allocation during nitrogen limitation, while xylitol related proteins were downregulated (in all cases adj. p-val < 0.01).

### 3.9 Protein changes under the reference condition are well correlated with the simulated fluxes

In response to nitrogen limitation, proteins involved in central nitrogen metabolism, such as glutamate dehydrogenase (GDH, RHTO_04650, RHTO_07718) and glutamine synthetase (GLN, RHTO_00673, RHTO_00401) were upregulated (Figure 3A). This response has been previously reported under nitrogen limitation for *R. toruloides* grown in both glucose and xylose (Zhu et al., 2012; Tiukova et al., 2019b). Additionally, activation of autophagy process has been described as a direct response via TOR activation to recycle nitrogenous compounds (Zhu et al., 2012; Tiukova et al., 2019b). Although initially expressed at a low level, upregulation of autophagy-related proteins (RHTO_05541, RHTO_06526) was detected.

**Figure 3.**
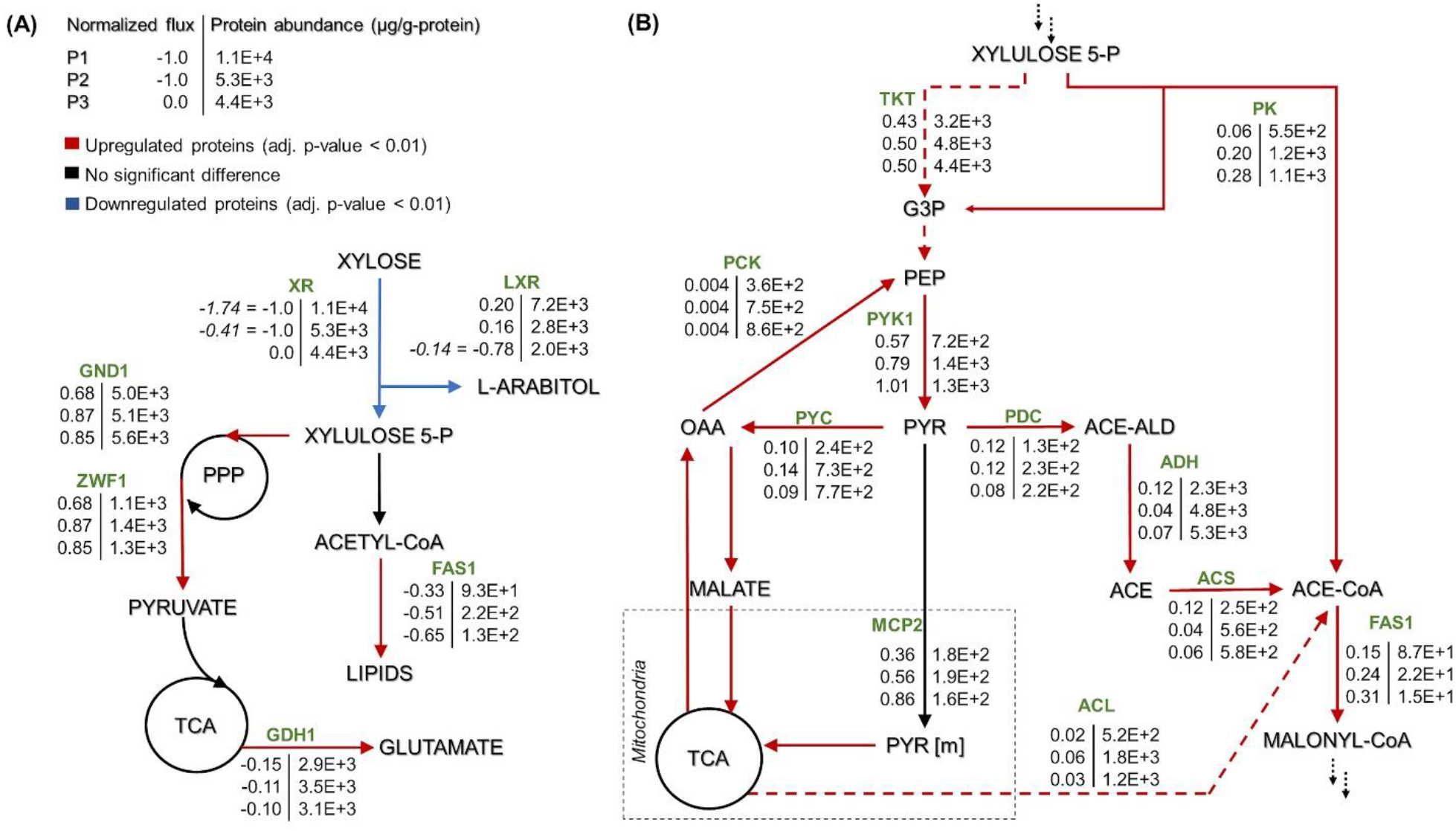
*R. toruloides* GEM-based fluxes (normalized to the substrate uptake) and measured protein abundances (μg/g-protein) under the three growth phases of reference cultivation on xylose (P1-P3). The main NADPH regeneration and utilization fluxes are presented with positive and negative values, respectively **(A)**. Representation of the central carbon metabolism illustrating xylulose 5-phosphate, pyruvate and acetyl-CoA branchpoints **(B)**. The fluxes were normalized by the substrate uptake rate in each phase (the absolute substrate uptake values (mmol/g_DCW_.h) are represented in italic in panel A (note that in P3, arabitol and xylitol were co-consumed)). The arrow colour indicates the significant protein allocation change under nitrogen limitation (P2 and P3) compared to nitrogen excess condition (P1); and the dashed arrow on panel B represent multiple reactions. All the fluxes and protein abundance are represented in the Supplementary Tables S3 and S7, respectively. G3P: glyceraldehyde 3-phosphate, PEP: phosphoenolpyruvate, PYR: pyruvate, OAA: oxaloacetate, ACE-ALD: acetaldehyde, TKT: transketolase, PK: phosphoketolase, PCK: phosphoenolpyruvate carboxykinase, PYK1: pyruvate kinase, PYC: pyruvate carboxylase, PDC: pyruvate decarboxylase, MCP2: mitochondria pyruvate carrier, ADH: alcohol dehydrogenase, ACS: acetyl-CoA synthase, ACL: ATP citrate lyase, and FAS1: fatty acid synthase. More abbreviations can be found in the Supplementary Table S4.

Proteins related to oxidative stress response showed upregulation for P2 and P3 (nitrogen limitation) compared to P1 (nitrogen excess). Catalase (CAT, RHTO_01370), which breaks down hydrogen peroxide in the peroxisomal matrix, was the most upregulated protein with a 6-fold increase under nitrogen limitation. Recent reports in oleaginous microorganisms showed that ROS is an important signalling molecules in response to various stresses (Shi et al., 2017). Nitrogen depletion is an example of such stress, leading to the accumulation of ROS (Liu et al., 2012; Fan et al., 2014; Chokshi et al., 2017) and higher activities of catalase and other antioxidant enzymes, suggesting that lipid accumulation under nitrogen depletion is mediated by oxidative stress (Yilancioglu et al., 2014).

The highest carbon fluxes detected with GEM analysis were further assessed to understand the level of their regulation. Xylose uptake decreased under the nitrogen limitation in P2 and was lacking in P3, which was also reflected in downregulation of proteins involved in xylose assimilation. Proteins belonging to arabitol metabolism, such as L-xylulose reductase (LXR, RHTO_00373) and D-arabinitol dehydrogenase (AD, RHTO_07844), were also downregulated in P2 and P3.

Approximately 4-fold increased transketolase (TKT, RHTO_03248) abundance compared to phosphoketolase (PK, RHTO_04463) was consistent with the simulated increased flux through the transketolase reaction. However, PK levels increased more than 2-fold under nitrogen limitation, which was consistent with the increased flux levels under the mentioned conditions. Furthermore, the magnitude of the PK increase in this condition was 50-60% higher than TKT (Figure 3B). Carbon was mainly channelled via TKT because it leans towards glycolysis and the oxidative branch of the PPP, which have been identified as the preferred pathway to regenerate NADPH (Lopes et al., 2020a, this study). GEM simulations revealed that xylose reductase (XR), fatty acid synthase and glutamate dehydrogenase consume most of the NADPH, which was regenerated by 6-phosphogluconate dehydrogenase (GND1), glucose-6-phosphate dehydrogenase (ZWF1), and L-xylulose reductase (LXR) (Figure 3A). With the exception of fatty acid synthase, these enzymes were also among the most abundant NADPH-dependent enzymes quantified.

In the oleaginous microorganism, cytosolic ATP-citrate lyase (ACL, RHTO_03915) is considered an important enzyme as a source of acetyl-CoA (Ratledge and Wynn, 2002; Koutinas and Papanikolaou, 2011). Under nitrogen limitation, this enzyme was found in higher levels compared to P1 (4- and 2-fold increase for P2 and P3, respectively). Cytosolic acetyl-CoA can also be supplied by acetyl-CoA synthase (ACS1, RHTO_08027), which was upregulated 2-fold in both P2 and P3, and by xylose metabolism via PK and phosphate acetyltransferase. However, FBA predicted that the majority of cytosolic acetyl-CoA originated from pyruvate via ACS1 and from PK and phosphate transacetylase (Figure 3B).

Acetyl-CoA can be transformed into malonyl-CoA, a substrate for FA synthesis, by the enzyme acetyl-CoA carboxylase (ACC1, RHTO_02004), which was 3.4- and 1.6-fold upregulated in P2 and P3, respectively. Then, FAs are produced by fatty acid synthases FAS1 (RHTO_02032) and FAS2 (RHTO_02139), which were significantly upregulated in the nitrogen-limiting phases. Such findings were in accordance with the predicted higher fluxes from acetyl-CoA towards FA synthesis in P2 and P3. Additionally, acetyl-CoA can enter the mevalonate pathway (MEV) through acetyl-CoA C-acetyltransferase (ERG10, RHTO_02048) to produce sterols and carotenoids. Despite the highest carotenoid content in P3, ERG10 and other proteins related to the MEV pathway, hydroxymethylglutaryl-CoA synthase (ERG13, RHTO_02305) and reductase (HMG1, RHTO_04045) were downregulated in P3. Proteins directly related to the carotenogenesis pathway were either not detected or did not show any significant difference. FBA predicted very low fluxes throughout carotenoid production.

### 3.10 Comparative proteomics at different growth phases under oxidative stresses

Although P2 of cultivation under light exposure (LIG_P2) showed higher carotenoid production rates than REF_P2, only two proteins (RHTO_06480 and RHTO_01160) were differentially expressed between those conditions (adj. p-value < 0.01). The functions of these proteins do not explain the higher production of carotenoids and lipids, suggesting that carotenoid production is regulated post-translationally.

Under oxidative stress, the adapted strain showed many similar changes to the reference condition while entering into nitrogen limitation but was also clearly differentiated as described by PC analysis (Figure 4B). Clustering of the significantly differentially expressed proteins was performed to understand the main changes compared to REF. Cluster showing significant differences among ADA and other conditions was enriched by the proteins from amino acid biosynthesis and the TCA cycle. Although TCA cycle was already upregulated under nitrogen-limiting conditions, more pronounced upregulation was detected in ADA, including the upregulation of citrate synthase (CIT1, RHTO_06406), malate dehydrogenase (MDH, RHTO_04363), and NADP^+^-dependent isocitrate dehydrogenase (IDH, RHTO_04315). NADPH regeneration by GND1 (RHTO_ 02788) in PPP was also upregulated. Similar to the REF under nitrogen limitation, FA biosynthesis was upregulated; however, β-oxidation, which is responsible for lipid degradation, remained lower and could at least partially explain the higher lipid accumulation under oxidative stress. Additionally, the carotenogenesis pathway was more activated since ERG10 and phytoene dehydrogenase (CRTI, RHTO_04602) were 2- and 4-fold upregulated, respectively (Figure 5A).

**Figure 4.**
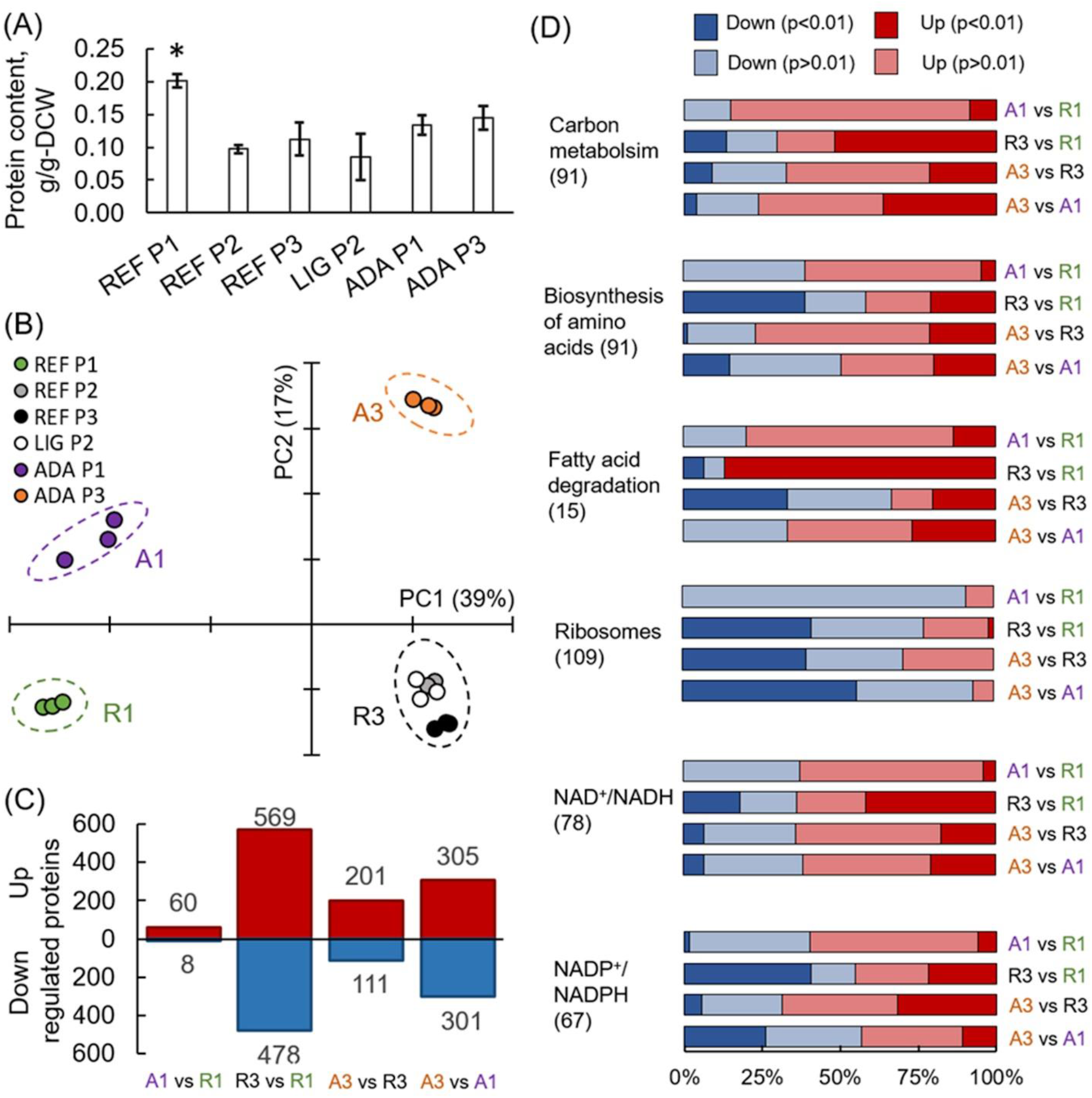
Proteomics results from *R. toruloides* studies on xylose under different environmental conditions (REF P1, P2, and P3; LIG P2; ADA P1 and P3). Total protein content in cells **(A)**. Principal component analysis **(B)**. Significantly (adj. p-value < 0.01) up- and downregulated proteins **(C)**. Gene set enrichment analysis based on the proteomics data, where GO groups were received from Uniprot, enzyme-metabolite interactions and subgroups from the rhtoGEM **(D)**. All previewed categories show significant difference at least in 2 comparisons (adj. p-value < 0.001). Number in brackets indicates proteins in each category. Panels **(C)** and **(D)** present comparisons between samples showing clear separation in PCA represented in (B).

**Figure 5.**
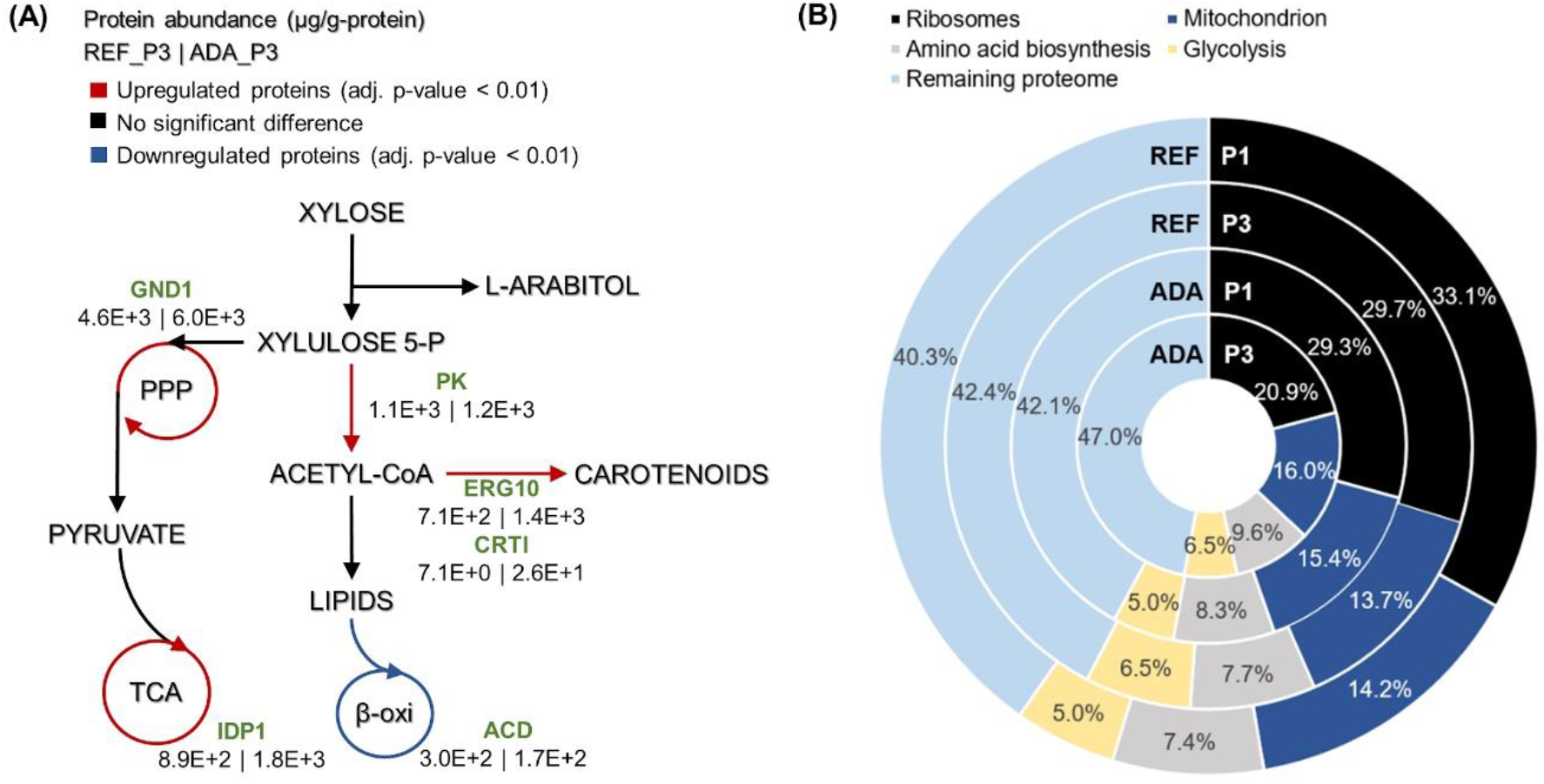
Main differences in protein allocation between the adapted strain under oxidative stress (ADA) and the parental strain under reference condition (REF) studied under the nitrogen limitation (P3). The arrow colour indicates the change in protein allocation in ADA compared to REF **(A)**. Proteome allocation into the most abundant metabolic groups for ADA and REF in P1 (nitrogen excess) and P3 (nitrogen limitation) **(B)**. GND1, 6-phosphogluconate dehydrogenase; IDP1, isocitrate dehydrogenase; PK, phophoketolase; ERG10, acetyl-CoA acetyltransferase; CRTI, phytoene dehydrogenase, ACD, acyl-CoA dehydrogenase.

As expected, enzymes involved in the oxidative stress response were upregulated in the ADA compared to REF; CAT exhibits a 16- and 5-fold increase for P1 and P3, respectively. For P3, other oxidative stress related proteins, such as S-(hydroxymethyl) glutathione dehydrogenase (RHTO_03559) and glutathione S-transferase (RHTO_00710), were also significantly upregulated (Supplementary Table S7).

### 3.11 Translation processes also play a crucial role in proteome allocation analysis

Absolute protein analysis allows comparisons of protein abundance levels between conditions and quantification of condition-dependent protein allocation patterns (Figure 5B). The top 100 of the most abundant proteins represented greater than 50% of the total proteome. Ribosomal proteins were the largest protein group, forming almost one-third of the total proteome in REF P1. However, these levels decreased to approximately 10% under the lower growth rate conditions in P3. For ADA, ribosomal protein allocation was already reduced to a lower level and decreased further, forming only 21% under the nutrient-limiting conditions in P3. As ribosomes are essential for achieving faster cell growth, the trade-off between allocation towards ribosomal proteins or energy generation pathways has been demonstrated previously (Nilsson and Nielsen, 2016; Sánchez et al., 2017, Kumar and Lahtvee, 2020). Interestingly, glycolysis was increased to the same extent under nitrogen-limiting conditions in the presence and absence of oxidative stress, but mitochondria and amino acid metabolism demonstrated significantly increased allocation for the ADA. The latter changes seemed to be responsible for the more efficient metabolism, providing higher biomass yields for adapted cells (Figure 5B and Table 1).

## 4 Discussion

Efficient microbial production of chemicals from sustainable resources is essential for the transition towards bioeconomy. *R. toruloides* has been considered as a potential microorganism to produce high-value products from biological resources, including hemicellulosic material, mainly composed of xylose. However, xylose metabolism in *R. toruloides* is still not completely understood. Only few studies have focussed on the metabolism of xylose assimilation in this oleaginous yeast (Jagtap and Rao, 2018; Tiukova et al., 2019b; Lopes et al., 2020a). Therefore, in our study, detailed physiology characterisation was combined with genome-scale modelling and absolute proteomics with the goal to investigate xylose metabolism in *R. toruloides* and use oxidative stress as a strategy to improve the production of lipids and carotenoids.

In this study, we demonstrated how *R. toruloides* growth on xylose exhibited three distinct phases, where most metabolic changes occurred after the transition into nitrogen limitation. Approximately 30% of consumed xylose by the parental strain accumulated into xylitol and arabitol during the first two growth phases probably to balance NADPH required for the growth or due to limitations in the abundance of xylulokinase. Although Jagtap and Rao (2018) reported that xylose conversion into D-arabitol by arabitol dehydrogenase regenerated one NAD^+^, the UniProt protein database for *R. toruloides* contains only sequences for LXR and for AD, lacking the levogyrous version of the latter. According to Fernandes and Murray (2010), fungi can produce both L- and D-arabitol. However, our proteomics analysis was able to detect only the enzymes for L-arabitol production, and the corresponding pathway was also used in our modelling approach (Supplementary Figure S1). Xylulokinase, an enzyme responsible for the last step in xylose metabolism, was not detected in our proteomic analysis, suggesting a low expression level that could limit xylose catabolism.

Under nitrogen-limiting conditions, increased flux via phosphoketolase was observed, corresponding well with enzyme upregulation. The phosphoketolase reaction saves carbon by producing an acetyl residue together with G3P directly from D-xylulose 5-P instead of requiring several reactions until pyruvate decarboxylation, potentially resulting in an increased product yield from substrate. However, cells seem to prefer the alternative pathway of transketolase as it provides more carbon towards glycolysis and oxidative PPP, where the majority of NADPH is regenerated. The overexpression of phosphoketolase and phosphotransacetylase enzymes in *Y. lipolytica* resulted in 20-50% increased lipid production (Niehus et al., 2018). Therefore, the upregulation of phosphoketolase could be partially related to the increased accumulation of lipids and carotenoids at the aforementioned phases. NADPH regeneration is an important mechanism for xylose assimilation and lipid synthesis. ME is considered a key enzyme in the recycling of NADPH for lipid biosynthesis (Ratledge and Wynn, 2002). However, according to our proteomic data and simulations, the majority of the reducing power is generated in the oxidative branch of the PPP. These results are consistent with previous 13C-labelling experiments with *Y. lipolytica* performed using glucose as the carbon source (Wasylenko et al., 2015). However, ME overexpression in *Rhodotorula glutinis* has led to a 2-fold increase in lipid accumulation (Li et al., 2013).

Induction of oxidative stress in *R. toruloides* cultivation was employed as a strategy to identify the metabolic changes that result in the higher levels of carotenoids and lipids. Surprisingly, under PER conditions, no changes in carotenoid production compared to REF were observed. However, the lipid content was two-fold increased, reaching a yield of 0.65 g/g_DCW_. The highest reported lipid production in *R. toruloides* is 0.675 g/g_DCW_ (Li et al., 2007) using a rich medium with glucose as a carbon source. The review by Shi et al. (2017) reports that different authors found evidence for ROS stress enhancing the formation of lipid droplets as an important signalling molecule in response to nitrogen starvation in oleaginous microorganisms. Cells under oxidative stress might possess an even greater supply of NADPH by channelling sugar catabolism to PPP to stabilise the redox balance and ROS clearance (Kuehne et al., 2015). Enzymes related to oxidative stress were upregulated not only in ADA but also in REF under nitrogen limitation (P2 and P3). Tiukova et al. (2019b) reported not only the upregulation of oxidative stress related enzymes during the lipid accumulation phases but also the upregulation of the beta-oxidation representing an ATP sink. According to Xu et al. (2017), lipid oxidation leads to accumulation of oxidative and aldehyde species in *Y. lipolytica*, reducing production performance. The same authors showed that overexpressing enzymes from the oxidative stress defence pathway resulted in industrially relevant lipid production (lipid titre of 72.7 g/L and content of 81.4%) by synchronising lipogenesis to cell growth and mitigating lipotoxicity.

The successive application of H_2_O_2_ in the cell through the ALE potentially overwhelmed the cellular antioxidant defence system, boosting carotenoid production. Increased accumulation of carotenoids and lipids was noted in all the phases of ADA compared to REF. In general, improved production can result from the upregulation of phosphoketolase, which led to a slightly more efficient production process. The mevalonate pathway was more activated by higher levels of ERG10, providing more precursors for carotenoid biosynthesis, which along with the upregulation of CRTI (enzyme of the carotenoid biosynthesis pathway) could explain the higher carotenoid levels. IDP1 and GND1 upregulation improved the capacity of NADPH regeneration. The upregulation of the oxidative stress defence pathway potentially diminished lipotoxicity. All the aforementioned factors combined with the downregulation of the β-oxidation could explain the higher lipid production by the adapted cells under oxidative stress (Figure 5A). No reasonable significant modifications in proteomics levels were observed under the light condition in contrast to the findings of Gong et al. (2019) for *R. glutinis.* However, irradiation increased final titres of carotenoids and lipids (75 and 40% improvement, respectively) likely due to post-translational mechanisms. Some fungi, such as *Mucor circinelloides*, harbour genes for carotenoid synthesis that are based on light-induced expression mechanisms (Quiles-Rosillo et al., 2005).

## 5 Conclusions

In this study, the detailed physiological characterisation of *R. toruloides* growth revealed that a considerable amount of xylose was converted into by-products, such as arabitol and xylitol. The accumulation of these by-products can be considered an overflow metabolism, contributing to the redox balancing during xylose catabolism. According to the simulations, the highest NADPH demand was related to substrate uptake and was about 5-fold higher compared with the NADPH levels required for lipid production. The main NADPH regeneration reactions were derived from the oxidative branch of PPP, and ME was underused.

Under nitrogen limitation, the parental strain showed some increased protein expression related to lipid synthesis (FAS complex) and oxidative stress response mechanisms. Interestingly, the adapted strain downregulated the FA degradation pathway, which combined with the upregulation of NADPH regeneration mechanisms and FA and carotenoid synthesis, led to better production performance. Additionally, this strain demonstrated increased allocation of proteins related to mitochondria and amino acid metabolism, potentially explaining the more efficient metabolism.

In general, a good correlation was noted between the predicted fluxes and determined protein abundances. Using data obtained in this study, we can design strategies of metabolic engineering to make the process economically viable by improving cell factory performance.

## 7 Author Contributions

MJP and NB share first co-authorship. MJP, NB, EAM and PJL designed the experiments. MJP performed the experiments. IB performed the metabolic flux simulations. MJP, NB, IB and PJL analysed the data. MJP, NB, IB, EAM and PJL wrote and revised the manuscript.

## 9 Acknowledgements

We thank the Proteomics Core Laboratory at University of Tartu for the proteome quantification.

