## Supplementary Figures for "Xylose Metabolism and the Effect of Oxidative Stress on Lipid and Carotenoid Production in *Rhodotorula toruloides*: Insights for Future Biorefinery"

**Figure S1.** Schematic representation of the pathway leading to the consumption of xylose by *R. toruloides*. All fluxes and complete legend can be found in the Supplementary Table S4.

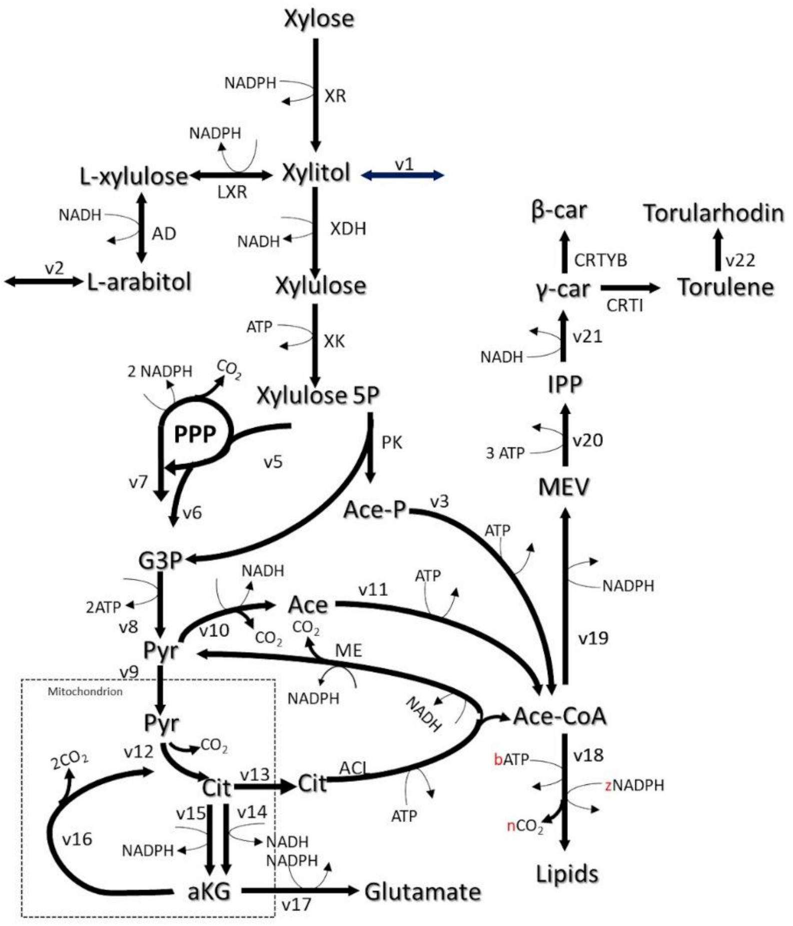

**Figure S2.** *R. toruloides* batch cultivation on xylose under light irradiation **(A)** and in presence of $H_2O_2$ **(B)**, and adapted strain under oxidative stress **(C)**. Dashed vertical lines define three observed growth phases. The specific growth rate (μ), biomass concentration, intracellularly accumulated lipid and carotenoid concentrations, extracellular metabolite profiles and $CO_2$ production profile in the outflow gas are presented. The values represent an average of three independent cultivation experiments; error bars represent standard deviation.

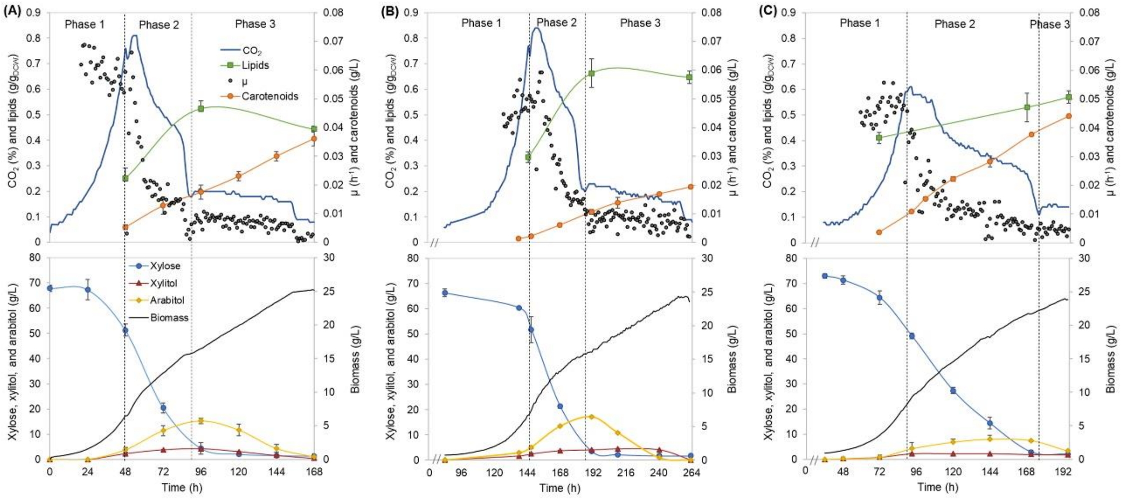


**Figure S3**. Changes in $\mu_{max}$ and lag phase time during the **(A)** 1st and **(B)** 2nd cycle of the adaptive laboratory evolution with $H_2O_2$. Error bars represent the standard deviation from three measurements.

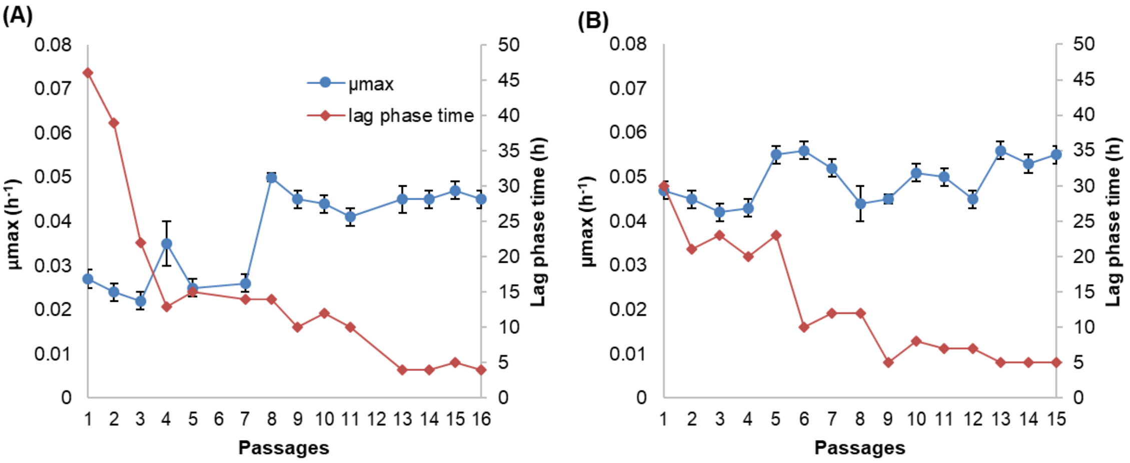

**Figure S4**. Carotenoids composition during the batch cultivation of *R. toruloides* under the studied conditions: reference condition (REF), under light irradiation (LIG) and in presence of hydrogen peroxide (PER), and the adapted strain under oxidative stress (ADA).

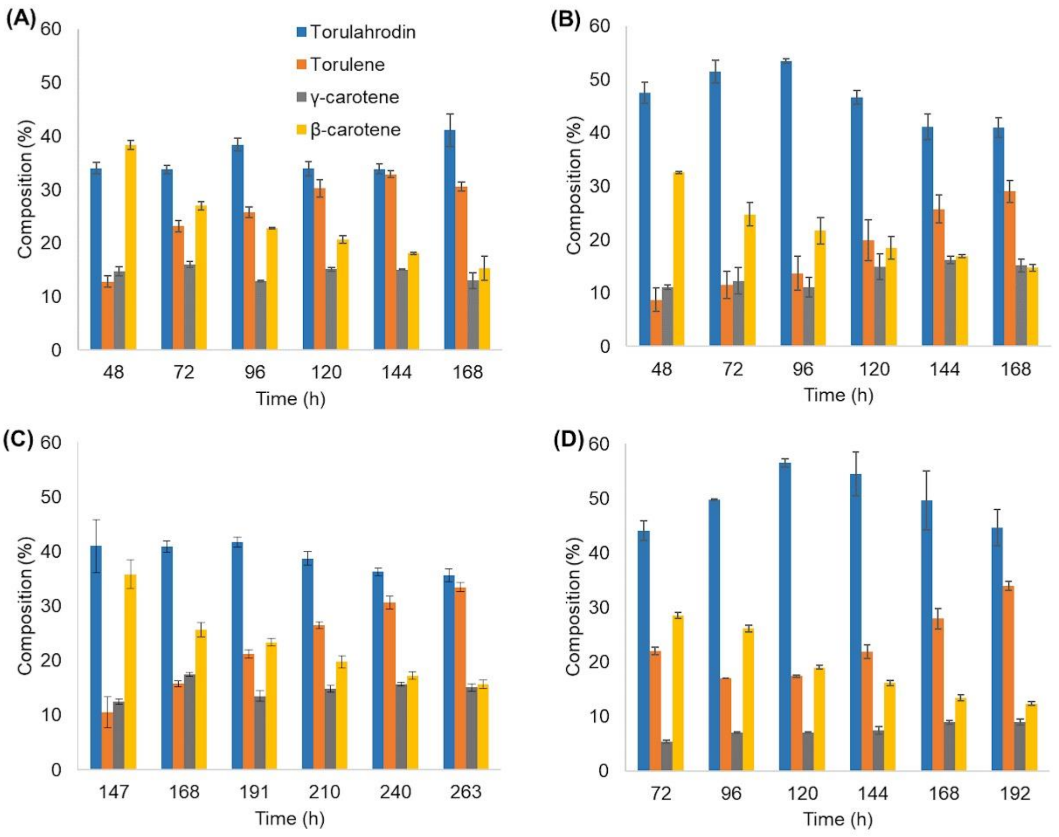
